# A marine fungus efficiently degrades polyethylene

**DOI:** 10.1101/2021.11.19.469330

**Authors:** Rongrong Gao, Rui Liu, Chaomin Sun

## Abstract

Plastics pollution has been a global concern. Huge quantities of polyethylene (PE), the most abundant and refractory plastic in the world, have been accumulating in the environment causing serious ecological problems. However, the paucity of microorganisms and enzymes that efficiently degrading PE seriously impedes the development of bio-products to eliminate this environmental pollution. Here, by screening hundreds of plastic waste-associated samples, we isolated a fungus (named *Alternaria* sp. FB1) that possessing a prominent capability of colonizing, degrading and utilizing PE. Strikingly, the molecular weight of PE film decreased 95% after the fungal treatment. Using GC-MS, we further clarified that a four-carbon product (named Diglycolamine) accounted for 93.28% of all degradation products after the treatment by strain FB1. We defined potential enzymes that involved in the degradation of PE through a transcriptomic method. The degradation capabilities of two representative enzymes including a laccase and a peroxidase were verified. Lastly, a complete biodegradation process of PE is proposed. Our study provides a compelling candidate for further investigation of degradation mechanisms and development of biodegradation products of PE.

## Introduction

Plastic deposition has accumulated tremendously due to its extensive production, widespread application and high resistance to biodegradation [1–3]. The global production has expanded to 464 million tons in 2018 [4], and 50% of them was discarded within a short period after use[4, 5]. Among the total accumulated plastic waste, polythene (PE) alone accounts for 64% [6, 7], and is considered as most ecological problematic due to its high molecular weight, strong hydrophobicity and highly inert chemically and biologically [8–12]. In this sense, it is an urgent need to find an efficient approach to degrade the PE waste for decontamination from the environment. The degradation of PE can occur by chemical, thermal, photo or biological degradation [13]. Recently, biodegradation using microorganisms has become a promising alternative for plastic recycling due to the mild and environmentally friendly reaction conditions required [14].

Among these microbes, fungi are potentially effective for the degradation of environmental PE [15–18]. Generally, fungi as well as other microbes are involved in the depolymerization, assimilation and mineralization processes of plastic degradation along with the involvement of a set of enzymatic system responsible for diverse steps including colonization, degradation, transformation and utilization [19–21]. The hydrophilic surfaces make the initial colonization on PE surface really difficult, however, microbial enzymes can efficiently promote the attachment of microorganisms to the surface of PE by improving the plastic hydrophilicity [22]. Once fungi colonize on the surface of PE, the degradation and utilization processes could be performed in combination with both intracellular and extracellular enzymatic systems, which enable them to use complex polymers as a source of carbon and electrons for further growth [16]. The extracellular enzymatic system consists of the hydrolytic system that formed mainly by nonspecific oxidoreductases, including versatile peroxidases, laccases, and unspecific peroxygenases [23]. The intracellular enzymatic system is normally mediated by the cytochrome P450 family epoxidases and transferases [24].

Thus far several fungi including *Aspergillus*, *Acremonium*, *Fusarium*, *Penicillium*, *Phanerochaete* [25], and corresponding enzymes including laccases, manganese peroxidase and lignin peroxidases [26], have been reported to potentially degrade PE. Although some progresses have been made about fungi-mediated PE degradation, the biocatalytic degradation of PE needs to be detailedly elucidated, especially the category and function of enzymes responsible for the whole degradation process. Further research is required on novel isolates from plastisphere ecosystems to deeply disclose the associated mechanisms about PE biodegradation process [16].

Here, using a large-scale screening approach, we isolated and defined a marine fungus, *Alternaria* sp. FB1, which could effectively colonize and degrade PE. Using various techniques, we further clarified the degradation effects and released products. Lastly, we discovered 153 potential enzymes associated with PE degradation via a transcriptomic approach, and verified the degradation effects of two representative enzymes. Lastly, we described the whole process of PE degradation and utilization mediated by strain FB1.

## Results and Discussion

### Discovery of a marine fungus that efficiently colonizes and degrades PE

To obtain microorganisms possessing capability of degrading PE from marine environments, about 500 sedimentary samples containing plastic contaminants were collected from different locations of Huiquan bay of Qingdao, China. Thereafter, these samples were respectively put into a flask filled with sterilized sea water supplemented with only commercial PE bag fragments as the sole nutrient source. After one-month incubation at 25 °C, in the flask incubated with the 6^th^ sample, obvious filament bundles attached to four corners of the PE film were observed, suggesting some fungi might be enriched from this sample. These filament bundles were thus removed from the PE film and observed through a light microscopy. Indeed, the filament bundles attached to the PE film showed typical characteristics of fungal hyphae. Thereafter, the filament bundles were cultured in the PDA medium for further purification. After several rounds of purification and ITS (Internal Transcribed Spacer) gene sequencing confirmation, one pure fungal strain named FB1 was isolated. The fungal strain FB1 showed high homology (higher than 99% identity) of ITS sequence with many *Alternaria* strains, which was further confirmed by phylogenetic tree analysis (Fig. S1). Considering no more physiological characteristics were investigated, the strain FB1 was designated as *Alternaria* sp. FB1 in this study. Consistent well with its capability of growing in the sea water supplemented with only PE bags, it is indeed capable of efficiently colonizing on the pure PE films after 3-day incubation with PE in the seawater (Fig. 1b). Moreover, the fungus could generate many substances similar to biofilm among different hyphae (Fig. 1c) and produce large amount of spores for further multiplication (Fig. 1d), strongly suggesting strain FB1 could utilize PE as a nutrient source for growth given the presence of very little organic matter in the sea water. Consistently, strain FB1 showed a much better growth status and stronger reproduction ability in the seawater supplemented with PE film than that in the seawater only (Figs. S2-S3), confirming its capability of using PE as a nutrient source for growth.

**Fig. 1.**
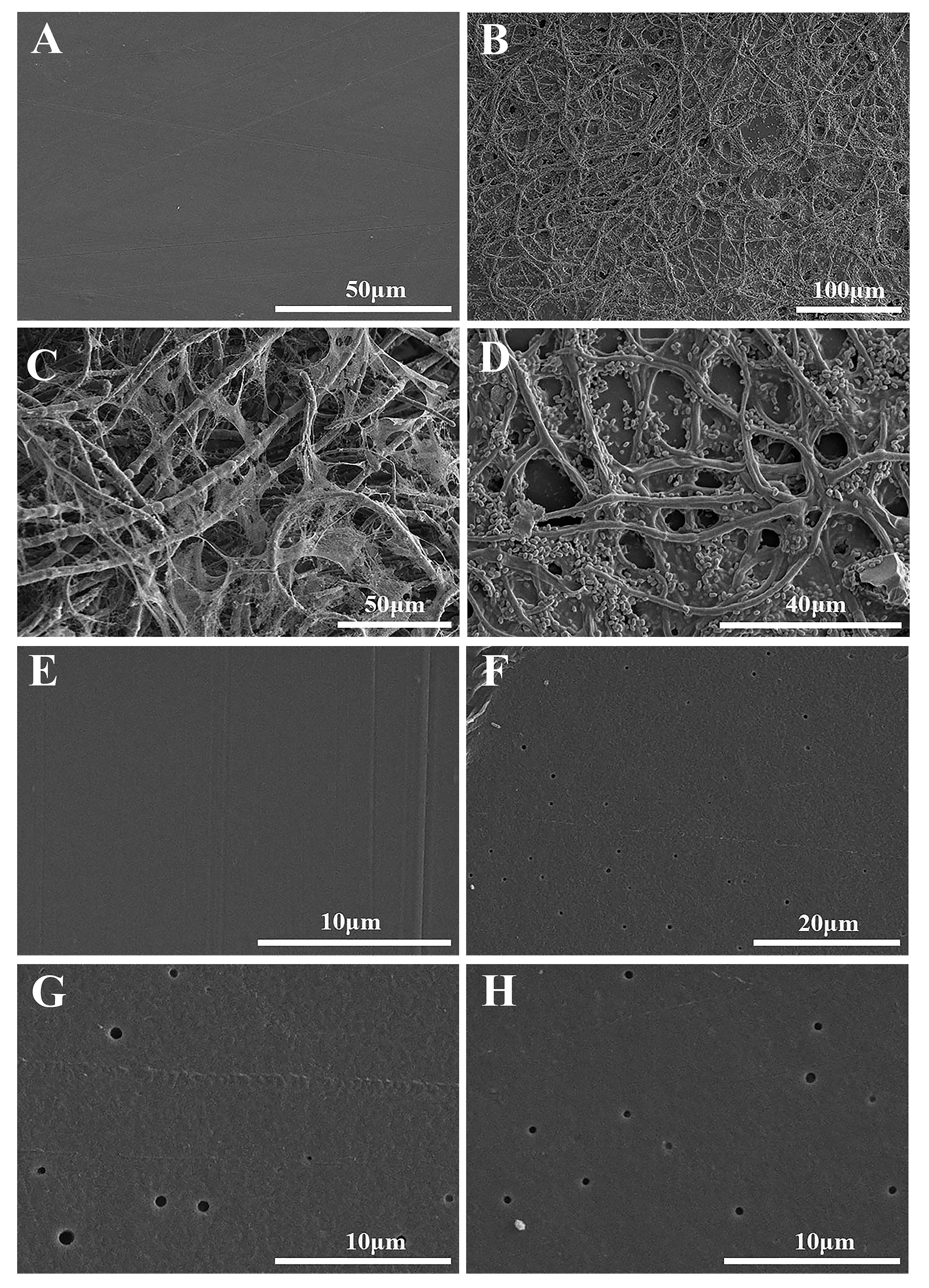
SEM observation of colonization and degradation effects of *Alternairia* sp. FB1 on the PE film. **a**, SEM observation of the PE film treated by the medium only for 7 days. **b-d**, SEM observation of the colonization of strain FB1 on the PE film after 7 days treatment. **e**, SEM observation of the PE film treated by the medium for 120 days. **f-h**, SEM observation of the degradation effects of strain FB1 on the PE film after 120 days treatment.

Next, after removing the microbial layer, we observed numerous holes in the PE surface (Fig. 1f), strongly indicating strain FB1 could effectively degrade PE films. The diameter of some holes could reach 0.5 μm, and some holes almost penetrated across the film (Figs. 1g-h). When we extended the treatment time to four months, the PE film could be thoroughly occupied by fungal hyphae and spores (Fig. 2c). The color of PE film changed from white to yellow and black, and the morphology of PE film became extreme curling and shrinking (Fig. 2c). On the other hand, while we only dropped some fungal cells at a special spot of the PE film and incubated in the sea water for about four months, almost all the growing area of fungus could separate from the original PE film (Figs. 2d-e). Taken together, we are confident that the marine fungus *Alternaria* sp. FB1 possesses prominent capabilities of colonizing, degrading PE and thereby utilizing as a nutrient source for growth.

**Fig. 2.**
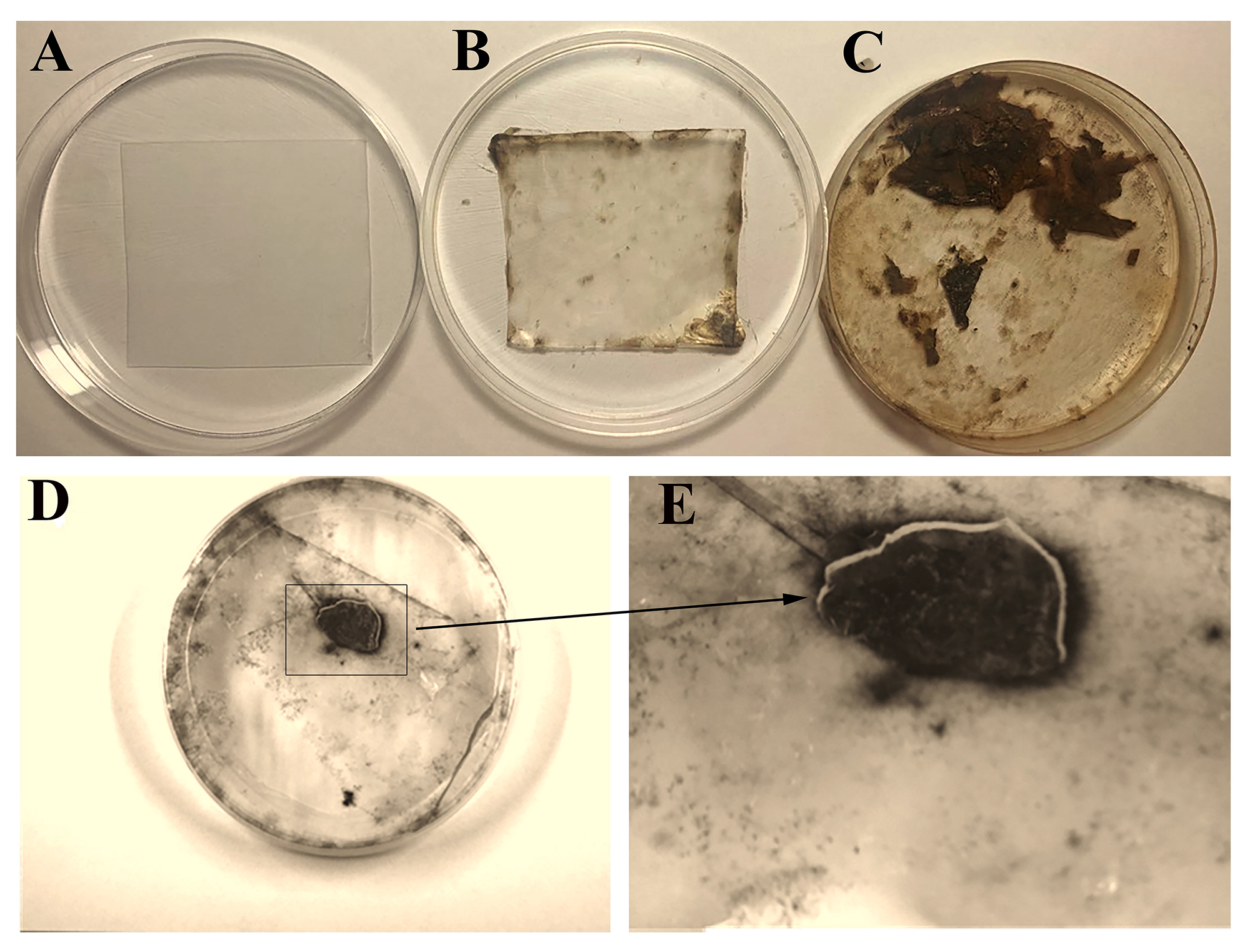
Significant morphological change of the PE film treated by *Alternairia* sp. FB1. **a**, Morphology of the PE film treated by the medium for 120 days. **b**, Morphology of the PE film treated by strain FB1 for 3 days. **c**, Morphology of the PE film treated by strain FB1 for 120 days. For panels a-c, the PE film was soaked in the medium without or with strain FB1. **d**, Morphology of the PE film treated by strain FB1 for 120 days. **e**, An amplifying observation of panel d. For panel d, the culture of strain FB1 was incubated in some area of the PE film. During the incubation course, proper amount of fresh medium was supplemented to maintain the growth of strain FB1.

Recently, it has been calculated that a range of 4.8-12.7 million tons of plastics enter the oceans annually [27], thus plastic pollutions have been recognized as the most common and durable marine contaminants. Consequently, the marine environment is becoming a hot spot to screen microorganisms possessing prominent plastic degradation capabilities [28], and our recent [29] and present works confirm the proposal. It is noting that *Alternaria* sp. FB1 is a typical representative of filamentous fungi that are found in different environments and some of them have evolved to adapt and grow even in terrestrial and marine environments under extreme conditions [16]. In particular, fungi are able to extend through substrates in their search for nutriments with their filamentous network structure, exploring and growing in places that are more difficult to reach for other microorganisms [16]. Indeed, our results clearly show that strain FB1 not only penetrates the PE film (Fig. 2) but also extends its growing location all over the surface of plastic (Fig. 1c), indicating fungi are good candidates for developing PE degradation bio-products.

### Verification of PE degradation effects conducted by *Alternaria* sp. FB1

Generally, several approaches are adopted to roughly evaluate visible changes in PE degradation, such as the formation of surface biofilms, holes, cracks, fragmentation, color changes, and surface roughness. The level of PE degradation can be further determined by SEM to verify the level of scission and attachment of the microorganisms, by Fourier Transform Infrared (FTIR) to analyze the microdestruction of the small samples, by X-Ray Diffraction (XRD) to evaluate the crystallinity degree, by Gel Permeation Chromatography (GPC) to estimate the depolymerization of PE long-chain structure [29, 30]. Next, we sought to further verify the PE degradation effects conducted by *Alternaria* sp. FB1 through above various approaches. First, FTIR analysis was conducted to detect the degradation effects. Compared to the control group, two extra FTIR spectra absorption peaks were observed in a 2-week fungus treated PE film (Fig. 3a, green curve). One absorption peak was observed in the vicinity of 1,715 cm^−1^, indicating the formation of carbonyl bonds (-C=O-), while the other absorption peak was observed at a wave number of 3,318 cm^−1^ and was attributed to hydroxyl groups. Moreover, the signal strength of above two peaks became much stronger when the treatment time was extended to four weeks (Fig. 3a, red curve). According to these key chemical bonds, we conclude that PE film treated by *Alternaria* sp. FB1 underwent major structural changes representing direct biodegradation by the fungus. Fungal treatment resulted in a cleavage of the PE polymer chain, which thereby reducing the molecular weight and increasing hydrophilicity of PE polymer.

**Fig. 3.**
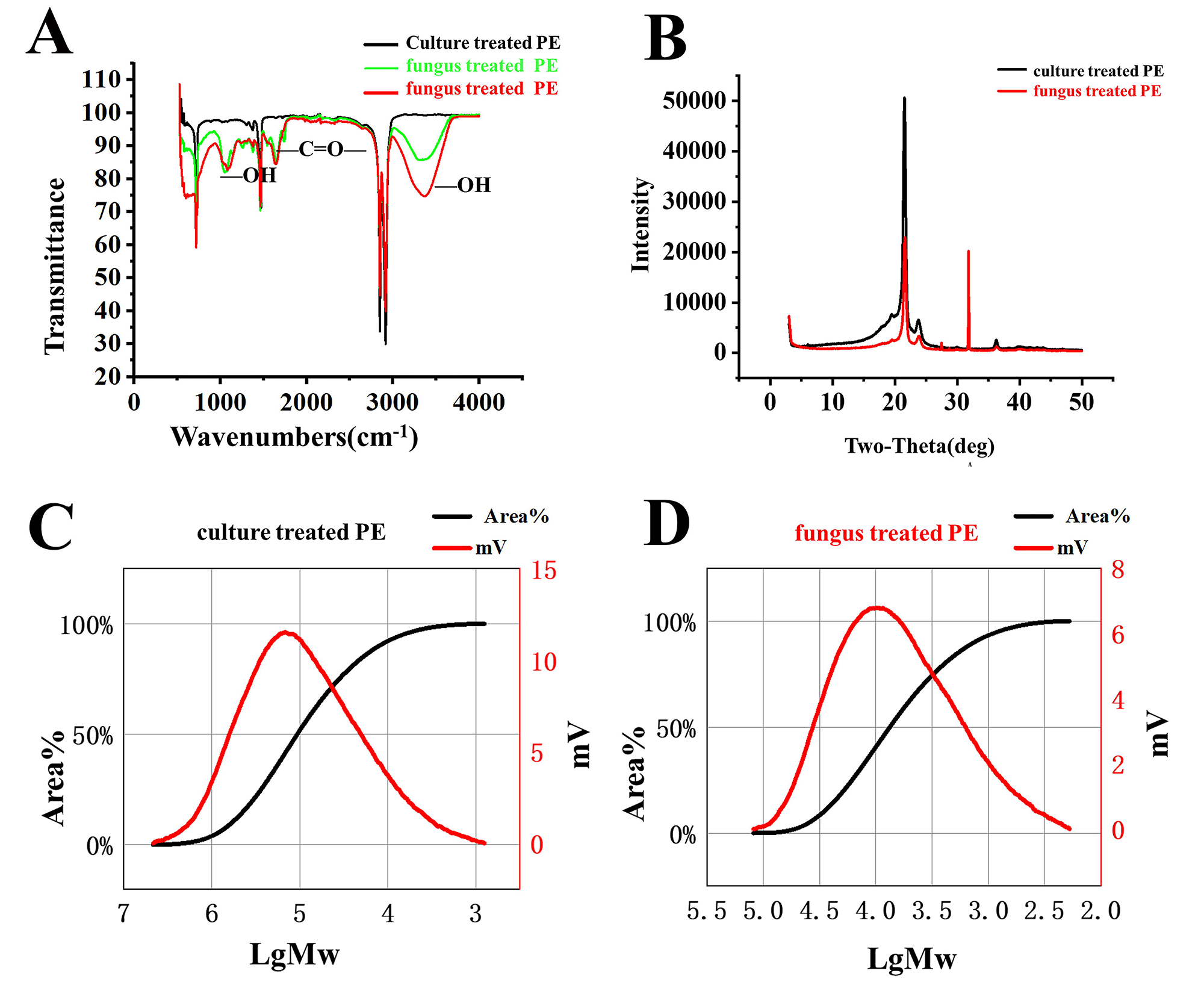
Verification of degradation effects of the PE film by *Alternairia* sp. FB1. **a**, FTIR analysis of the PE film treated by the medium without or with strain FB1 for two and four weeks. **b**, XRD analysis of the PE film treated by the medium without or with strain FB1 for four weeks. **c, d** GPC analysis of the PE film treated by the medium without (c) or with (d) strain FB1 for120 days.

On the other hand, through XRD analysis, we found that PE film treated by strain FB1 for 28 day s showed an evident reduced relative crystallinity degree, as measured by peak-differentiating and imitating calculations, resulting in a decrease from 62.79% to 52.02% (Fig. 3b). The XRD results clearly indicate that fungal treatment could significantly change the structure of molecular arrangement of PE polymer. Lastly, GPC was performed to determine the number-average molecular weight (Mn), molecular weight (Mw) and molecular weight distribution (MWD) of fungus treated PE films, which are three key indicators of the scission and degradation of plastics. After a 120-day treatment, the Mns of fungus-treated PE and medium-treated PE were respective 3,223 and 29,218, leading to a 9-fold decrease; the Mws of fungus-treated PE and culture-treated PE were respective 1,1959 and 231,017, resulting in a 20-fold decrease. Consistently, the MWD of fungus-treated PE (Fig. 3d) showed a markedly decrease trend compared to the culture-treated PE (Fig. 3c). The decrease of MWD and increase of the proportions of lower molecular weight fragments strongly suggested the occurence of depolymerization of the PE long-chain structure.

Through above techniques, many fungal strains belonging to general *Aspergillus*, *Penicillium* as well as *Fusarium* were found to be potentially efficient for PE degradation based on weight loss, molecular weight decrease and reduction in tensile strength [18]. In contrast, only one fungus belonging to the genus *Alternaria* was reported to cooperate with other fungi within a consortium to degrade the PE film [31]. Actually, the genus *Alternaria* includes more than 250 species and is ubiquitously distributed in diverse terrestrial and marine environments [32]. Our study clearly shows that *Alternaria* sp. FB1 possesses a prominent capability of degrading PE film: the molecular weight of PE film could be decreased 95% after fungal treatment, indicating this fungal strain as well as other *Alternaria* members has great potentials to develop plastic degradation products.

### Analysis of PE degradation products by Gas Chromatography-Mass Spectrometer (GC-MS) analysis

To further explore the details of PE degradation conducted by strain *Alternaria* sp. FB1, the degradation products were analyzed by GC-MS. For the 60-day treatment sample, the major retention time peaks corresponding to 17.58 min, 16.70 min, 18.98 min, 15.69 min and 17.16 min are the top 5 based on area percent calculation (Fig. S4). In contrast, for the 120-day treatment sample, only one predominant retention time peak corresponding to 7.75 min was shown, accounting about 93.28% of all peaks’ area (Fig. S5). Next, constituents existing in the above six retention time peaks were further identified by MS. The results revealed that within the 60-day treated sample the carbon numbers of each product ranged from 12 to 30 (Fig. 4a, Supplementary Table S1), and the product (1-monolinoleoylglycerol trimethylsilyl ether) possessing 27 carbons was predominant, accounting for 51.24% of all products (Fig. 4c). The rest predominant products were hexanedioic acid bis(2-ethylhexyl) ester (16.42%), squalene (13.89%), tributyl phosphate (7.1%), cycloheptasiloxane tetradecamethyl-(3.45%), cyclohexanamine N-cyclohexyl-(2.33%), 13-Docosenoic acid methyl ester, (Z)-(7.9%) (Fig. 4a and Supplementary Table S1). In contrast, in the 120-day treated sample, the carbon number of corresponding products ranged from 3 to 27 (Fig. 4b and Supplementary Table S2), and the product (Diglycolamine) possessing 4 carbons was the most predominant one, accounting for 93.28% of all products (Fig. 4c). Obviously, the proportion of product possessing smaller molecular weight significantly increased along with the extension of treatment time from 60 to 120 days, strongly suggesting that more evident degradation occurred after 120-day treatment by strain FB1.

**Fig. 4.**
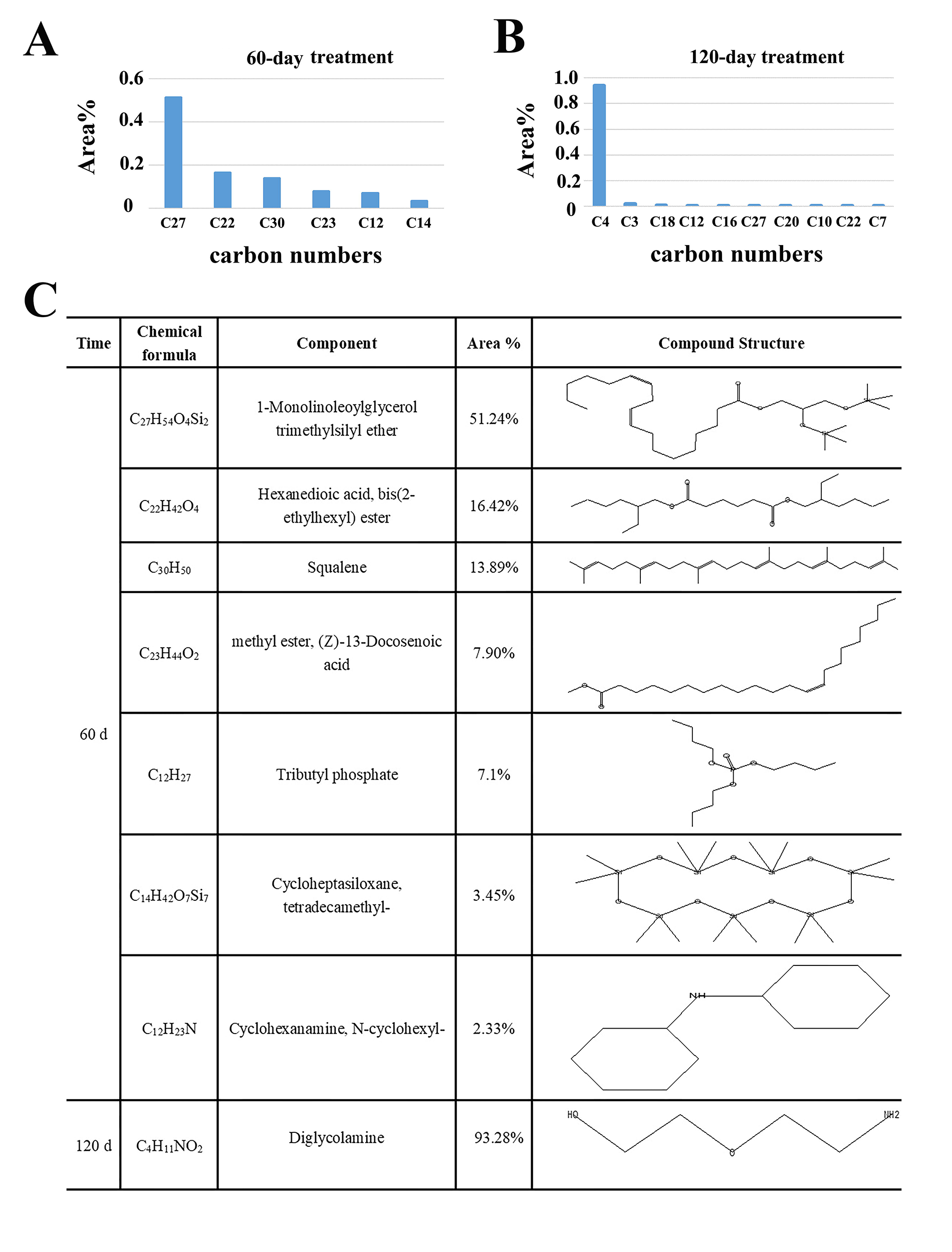
GC-MS analysis of products released from the PE film treated by *Alternairia* sp. FB1. **a**, The carbon number and respective proportion of degradation products released from the PE film treated by strain FB1 for 60 days. **b**, The carbon number and respective proportion of degradation products released from the PE film treated by strain FB1 for 120 days. **c**, The chemical formula, component, proportion and chemical structure of major products released from the PE film treated by strain FB1 for 60 days and 120 days.

Although some fungal strains have been identified as candidates for PE degradation[18], the degradation products are yet obscure. Notably, after 120 d treatment by strain FB1, the predominant degradation product is identified as Diglycolamine that possessing only four carbons (Fig. 4c). Diglycolamine, one of the alkanolamine solvents, produces total organic acid anions as degraded products[33, 34], which might contribute to the energy metabolism by some unknown pathway. We are confident that Diglycolamine is not a fungal metabolic product based on different database searches, however, it is still not clear how does Diglycolamine derive from the PE long chain and whether it will be degraded further or directly utilized by the fungus. Nevertheless, our study provides a hint for researchers to explore the degradation products of PE in the future.

### Transcriptomic profiling of the plastic degradation process

These parameters obtained from the FTIR, XRD, GPC as well as GC-MS can be used as indicators of microbial action, however, these results do not reflect the metabolic responses of microorganisms. To explore the plastic degradation process and potential mechanisms mediated by strain FB1, we performed a transcriptome analysis of this fungus cultured in the medium supplemented either with or without PE for 45 days. Combined with our gene expression analyses, we discovered 153 potential enzymes closely associated with biodegradation and the expressions of their encoding genes were significantly upregulated (Fig. 5a). In summary, these enzymes include 3 peroxidases, 3 laccases, 26 hydroxylases (4 hydroxylases, 15 monooxygenases, 7 oxygenases), 49 dehydrogenases, 18 oxidoreductases, 10 oxidases, 22 reductases, 16 esterases, 4 lipases and 2 cutinases. In particular, the transcription levels of laccase encoding gene (Gene id: evm.TU.contig_8.535), peroxidase encoding gene (Gene id: evm.TU.contig_5.872) and oxidoreductase encoding gene (Gene id: evm.TU.contig_5.292) were respectively increased about 23, 44 and 102 folds when compared the expressions of these genes under conditions supplemented with or without PE, strongly suggesting the key role s of these enzymes in the process of PE-degradation mediated by strain FB1.

**Fig. 5.**
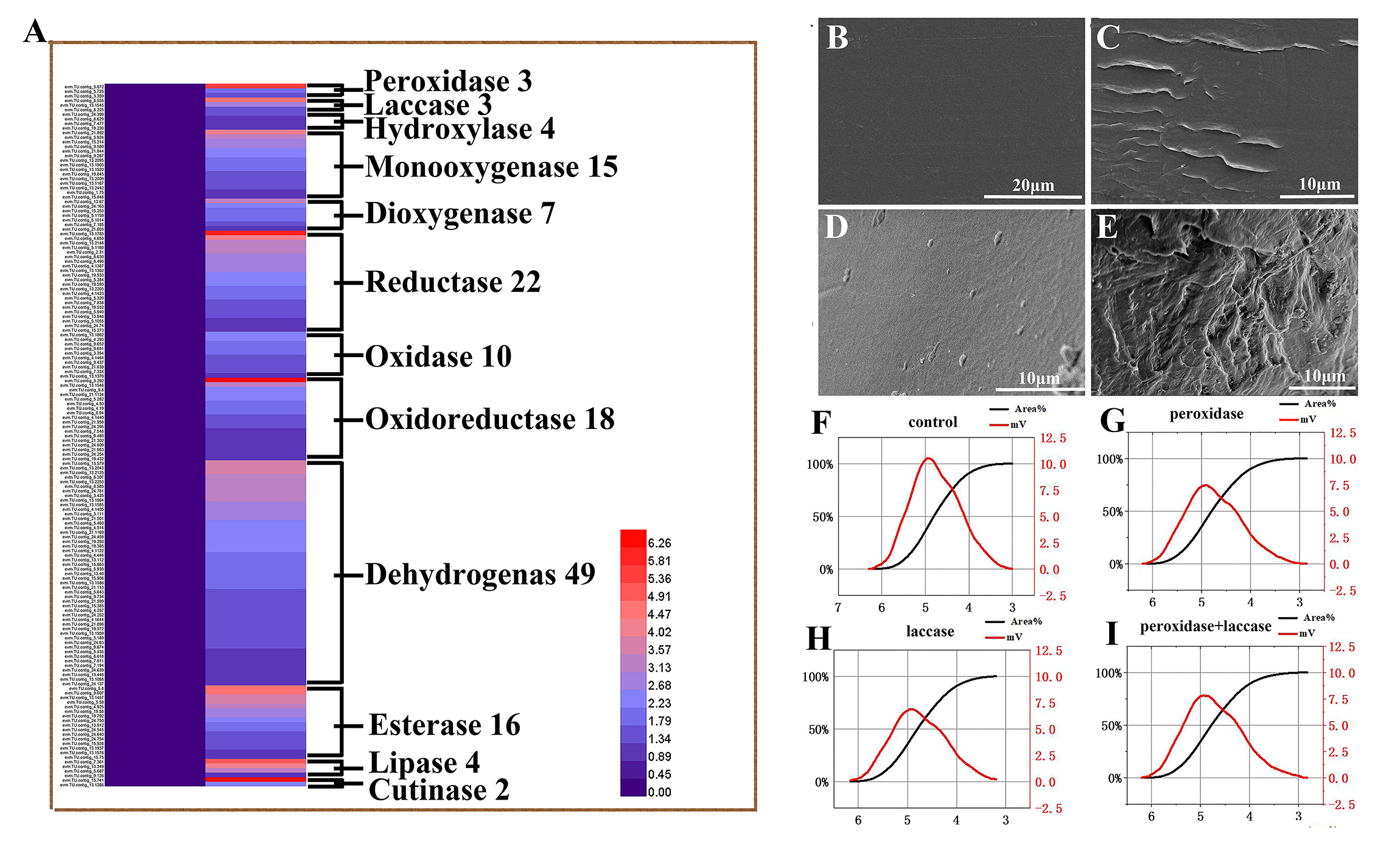
Transcriptomic analyses of PE degradation processes directed by *Alternairia* sp. FB1. **a**, A heat map showing the significantly up-regulated genes encoding enzymes with potential PE-degradation activities in strain FB1 that incubated with the PE film for 45 days. The number after corresponding enzymes’ name indicates the amount of genes whose expressions were significantly up-regulated (cutoff >2 folds). **b**, SEM observation of the PE film treated by the sterilized seawater for 48 h. **c**, SEM observation of the PE film treated by 0.1 mg/mL glutathione peroxidase (evm.model.contig_3.359) at 30 °C for 48 h. **d**, SEM observation of the PE film treated by 0.1 mg/mL laccase (evm.model.contig_8.535) at 30 °C for 48 h. **e**, SEM observation of the PE film treated by both 0.1 mg/mL glutathione peroxidase (evm.model.contig_3.359) and 0.1 mg/mL laccase (evm.model.contig_8.535) at 30 °C for 48 h. **f**, GPC analysis of the PE film treated by the sterilized seawater for 48 h. **g**, GPC analysis of the PE film treated by 0.1 mg/mL glutathione peroxidase (evm.model.contig_3.359) at 30 °C for 48 h. **h**, GPC analysis of the PE film treated by 0.1 mg/mL laccase (evm.model.contig_8.535) at 30 °C for 48 h. **i**, GPC analysis of the PE film treated by both 0.1 mg/mL glutathione peroxidase (evm.model.contig_3.359) and 0.1 mg/mL laccase (evm.model.contig_8.535) at 30 °C for 48 h.

To further verify the transcriptomic results, we overexpressed two putative PE degrading enzymes including glutathione peroxidase (evm.model.contig_3.359) and laccase (evm.model.contig_8.535) in *E. coli* cells (Fig. S26), and checked their respective degradation effects on PE films in 48 h. Notably, these two enzymes showed degradation effects on PE films compared to the control (treated by sterile seawater, Fig. 5b), obvious cracks and signs of plastic film degradation were observed by the SEM (Figs. 5c-d). Especially, glutathione peroxidase and laccase showed a clear synergetic degradation effect on the PE film (Fig. 5e). Consistently, the GPC analyses toward to both Mn and Mw of PE films treated by above two enzymes alone or together showed similar patterns to those of SEM observations. That is, the combined utilization of two enzymes led to a much higher degradation rate than those of single enzyme (Figs. 5f-i). The Mn and Mw of the PE film treated by both glutathione peroxidase and laccase were respective 20904 and 109202, which showed about 18% and 7% (Fig. 5i) decreases compared to those of control (Mn and Mw are 25516 and 116240, respectively, Fig. 5f). Future studies are required to test the degradation effects of more enzymes revealed by the transcriptomic results and develop a combined enzyme system for highly effective degradation of PE. Taken together, these knowledges greatly facilitate the protein and strain engineering for enhanced PE degradation performance, to meet the requirements for future industrial applications.

### A proposed model of biodegradation process of PE

Based on the combination of our genomic and transcriptomic data as well as previous reports, we propose a detail PE-degradation process (Fig. 6). Briefly, the process of PE biodegradation can be divided into four stages: colonization/corrosion, depolymerization, assimilation and mineralization [25, 35]. In the colonization stage, individual species or microbial consortium form a biofilm attached on the PE surface [25]. Due to the interaction with the various extracellular enzyme produced by microorganisms, the polymer surface was deteriorated and its hydrophobicity undermined. Then the long chain of the polymeric structure was broken down and was cut into small fragment by the action of a series enzymes secreted by the fungus. The initial and rate-determining step is the oxidation of PE by some oxidative enzymes such as peroxidase, oxygenase and laccase [15, 36, 37], which leading to a reduction of molecular weight. After the oxidation, the PE polymer is destructed, the molecular weight decreases, and carbonyl groups are introduced along the polyethylene chain. The decrease of molecular weight enables transport of PE small chain molecules through the cell membrane, it also makes degradation intermediates easier to be recognized and attacked by fungal enzymatic systems such as hydroxylase, monooxygenase, oxygenase, dehydrogenases, oxidoreductases esterases, lipases as well as cutinases. Given the chemical similarity between PE and alkanes, it has been suggested that the metabolic pathways for degradation of alkanes and PE are highly similar once the size of PE molecules decrease to an acceptable range for enzyme attack [25, 38]. In this sense, the PE degradation intermediates are further catalyzed by terminal oxidation monooxygenase to alcohol, which is further oxidized by alcohol and aldehyde dehydrogenases [39], and the resulting fatty acids enter the β-oxidation cycle. In parallel, the PE degradation intermediates are also catalyzed by sub-terminal oxidation monooxygenase to secondary alcohols, which are oxidized to ketones by alcohol dehydrogenase. A Baeyer-Villiger monooxygenase converts ketones to esters, which are subsequently cleaved by an esterase, cutinase and lipase. This leads to the formation of fatty acids and then degraded by β-oxidation. In the assimilation process, some small water-soluble intermediates with short chains produced by depolymerization are recognized by the receptors and then transported across the membrane into the microorganism, and thereby participating in a variety of metabolic activities and contributing to the cell growth. Finally, some metabolites and the non-assimilated products generated in the assimilation process are completely absorpted and utilized in mineralization, and are further converted to energy, carbon source as well as the CO_2_ and H_2_O.

**Fig. 6.**
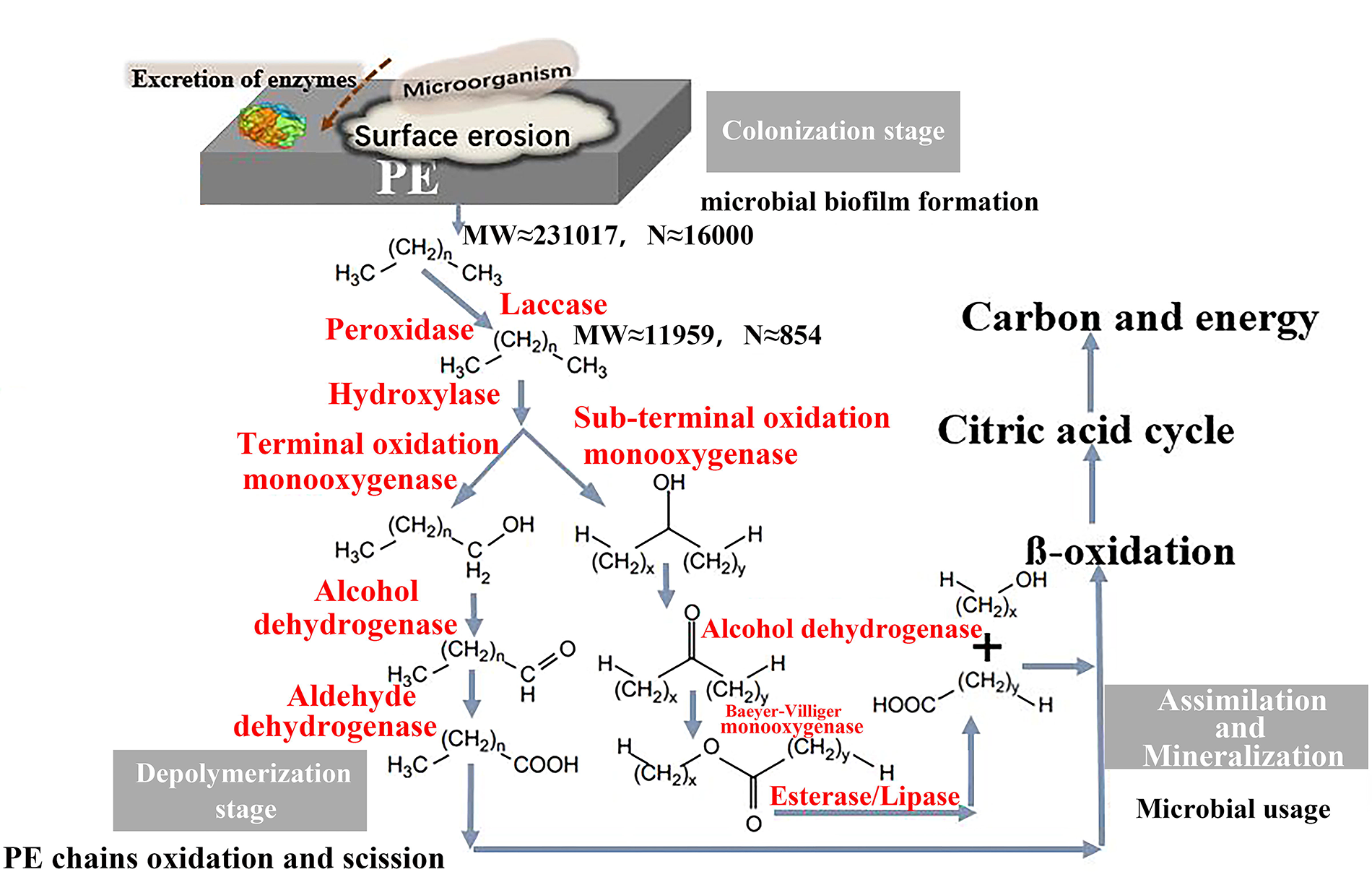
A proposed PE biodegradation model by the fungus. There are four stages in the PE biodegradation process: colonization/erosion, depolymerization, assimilation and mineralization. In this process, the PE polymer was degraded into small fragments step by step, then finally was converted to energy, carbon source as well as the CO_2_ and H_2_O. The detailed description of this model was shown in the results part.

## Conclusions

In our present study, we successfully obtained a marine fungus, *Alternaria* sp. FB1, which can efficiently colonize and degrade PE through forming numerous holes that across the film. Through SEM, FTIR and XRD approaches, we systematically verified the typical degradation indications including colonization, scission as well as microdestruction of PE film treated by strain FB1. Using GPC assay, we estimated the depolymerization of PE long-chain structure and find the molecular weight of PE film was decreased 95% after fungal treatment. Using GC-MS, we further clarified that a four-carbon product Diglycolamine was the most predominant (accounting for 93.28% of all products) degradation product after 120 days treatment by strain FB1. We defined the responses of this fungus directing plastic degradation through a transcriptomic method, showing the expressions of genes encoding 153 potential enzymes (including 3 peroxidases, 3 laccases, 26 hydroxylases, 49 dehydrogenases, 18 oxidoreductases, 10 oxidases, 22 reductases, 16 esterases, 4 lipases and 2 cutinases) are significantly up-regulated. The degradation effects of two representative enzymes revealed by the transcriptomic method were further verified by both SEM and GPC approaches. Lastly, three potential steps (including colonization, depolymerization and assimilation/mineralization) involved in biodegradation of PE are proposed.

## Methods

### PE plastics used for different assays

Three kinds of PE plastic are used in this study, including commercial PE bags, type ET311350 PE plastic (0.25 mm in thickness) and type ET311126 PE plastic (0.025 mm in thickness), the latter two are additive-free plastic films and are purchased from the Good Fellow Company (UK). All PE films are treated with 75% ethanol and air-dried in a laminar-flow clean bench prior to use.

### Screening, isolation and identification of marine microorganisms capable of degrading PE

To screen marine microorganisms capable of degrading PE, roughly 500 plastic debris samples were collected from the intertidal locations in the Huiquan Bay (Qingdao, China), and kept in flasks supplemented with filtered sea water and commercial PE films at room temperature (about 25 °C) for different periods. During this course, the films were checked by eyes and those covered by bacterial biofilm or fungal hyphae were observed by light microscopy to confirm the colonization of microorganisms. PDA medium (potato 200 g, glucose 20 g, agar 15~20 g, distilled water 1000 mL, natural pH) was used to purify the fugal strain FB1, and its purity was confirmed by PCR with the primers (ITSF: 5’-TCCGTAGGTGAACCTGCGG-3’; ITSR: 5’-TCCTCCGCTTATTGATATGC-3’) for identifying fungal ITS sequence. After the fungus was purified, the minimal medium (0.005 g yeast extract, 0.01 g peptone, 0.002 g xylose in 1 L filtered and sterile seawater, pH 7.0) was utilized for all growth and degradation assays of strain FB1 if not specified.

### Microscopic observation

The morphology and colonization of fungus on the PE film were observed and photographed by an inverted microscope (NIKON TS100, Tokyo, Japan) or scanning electron microscope (Hitachi S-3400N, Japan). The fungus or PE films were routinely observed by the inverted microscope according to the instruction. To observe the colonization of fungus on the plastic, PE films treated by medium or strain FB1 were soaked in 5% glutaraldehyde for cell fixation and were then dehydrated with 30%, 50%, 70%, 90%, 100% graded ethanol for 10 min each and critical-point-dried with CO_2_. Dried specimens were sputter coated for 5 min with gold and platinum (10 nm) using a Hitachi MC1000 Ion Sputter (Japan). To observe the plastic degradation effects, medium- or fungus treated PE films were washed in ultrasonic cleaner with 3% H_2_O_2_, 75% ethanol, and then distilled water to remove the biofilms thoroughly [40]. And the observation by SEM was performed as described above.

### Fourier Transform Infrared (FTIR) analysis

For FTIR analysis, PE films were recovered after a 2-week or 4-week fungus- or medium treatment. Films were then successively rinsed in ultrasonic cleaner with 1% SDS, distilled water, and then 75% ethanol [41, 42]. After air drying, PE films were recorded over the wavelength range of 450-4000 cm^−1^ at a resolution of 1 cm^−1^ using a Nicolet-360 FTIR (Waltham, USA) spectrometer operating in ATR mode [43]. Thirty two scans were taken for each spectrum.

### X-Ray Diffraction (XRD) analysis

XRD was performed using a Bruker D8 Advance instrument with a wavelength of 1.5406 angstrom of CuKα ray. The XRD tube current was set as 40 mA, and the tube voltage was set as 40 kV. Measurements for PE were set in the angle range from 2θ = 3° to 2θ = 50° at a rate of 1°/min [44].

### Gel Permeation Chromatography (GPC) analysis

The molecular weight of PE films treated by medium or fungus was determined by GPC on an Agilent PL-GPC220 (Agilent Technologies, USA) equipped with Agilent PLgel Olexis 300 × 7.5 mm columns and operating at 150 °C [45]. Trichlorobenzene was used as a mobile phase (1 mL/min) after calibration with polystyrene standards of known molecular mass. A sample concentration of 1 mg/mL was used [43, 46].

### Gas chromatography-mass spectrometry (GC-MS) analysis

The products of PE biodegradation were detected by GC-MS. Briefly, after 60-day or 120-day incubation of strain FB1 in the minimum medium supplemented with or without PE, corresponding cell suspension was centrifuged (12,000 *g*, 30 min, 4 °C) to collect the supernatant. The supernatant was freeze-dried and re-dissolved in 1 mL dichloromethane, then 2 μL filtered supernatant was used for GC-MS analysis performed on TRACE_1300GC-ISQ_LT GC-MS system (Shismadzu, Japan) equipped with a TG-5ms (30 m long, 0.25 mm internal diameter and 0.25 μm thickness) chromatographic column [47]. The injection-port was set at 300 ºC. During operation the column temperature was held for 4 min at 50 °C, then raised to 300 °C at 20 ºC rise per min, and finally, held for 15min at 300 °C. The flow rate was 0.800 mL/min. Helium was used as a carrier gas. Ions/fragments were monitored in scanning mode through 30-450 Amu.

### Genomic and transcriptomic analyses

To sequence the genome of strain FB1, the fungus was cultured in the PDA medium for 5 days, then the cells were collected and total DNAs were extracted with a DNeasy Blood and Tissue Kit (Qiagen, Germany) according to the instructions. Genomic sequencing was performed by Novogene (Tianjin, China) [48]. Sequencing libraries were generated using NEBNext® Ultra™ DNA Library Prep Kit for Illumina (NEB, USA) following manufacturer’s recommendations and index codes were added to attribute sequences to each sample [49]. Five databases were used to predict gene functions, including GO [50], KEGG [51], COG [52], NR [53] and Swiss-Prot [54].

For the transcriptomic analysis, strain FB1 was cultured in the minimum medium supplemented with or without PE films (type ET311350) for 45 d. Thereafter, the fungal cells were collected for further transcriptomic analyses performed by Novogene (Tianjin, China). Briefly, total RNAs from each sample were extracted and RNA degradation and contamination were monitored on 1% agarose gels. RNA purity was checked using the NanoPhotometer® spectrophotometer (Implen, USA). RNA concentration was measured using Qubit® RNA Assay Kit in Qubit® 2.0 Flurometer (Life Technologies, USA). RNA integrity was assessed using the RNA Nano 6,000 Assay Kit of the Bioanalyzer 2100 system (Agilent Technologies, USA). A total amount of 1 μg RNA per sample was used as input material for the RNA sample preparations. Sequencing libraries were generated using NEBNext® Ultra™ RNA Library Prep Kit for Illumina® (NEB, USA) following manufacturer’s recommendations and index codes were added to attribute sequences to each sample. The clustering of the index-coded samples was performed on a cBot Cluster Generation. After cluster generation, the library preparations were sequenced on an Illumina Hiseq platform and 125 bp/150 bp paired-end reads were generated. Raw data (raw reads) of fastq format were firstly processed through in-house perl scripts [55]. Reference genome and gene model annotation files were downloaded from genome website directly. Index of the reference genome was built using Hisat2 v2.0.4 and paired end clean reads were aligned to the reference genome using Hisat2 v2.0.4. HTSeq v0.9.1 was used to count the reads numbers mapped to each gene. And then FPKM of each gene was calculated based on the length of the gene and reads count mapped to this gene. Differential expression analysis was performed by using the DESeq R package (1.18.0)[56]. Gene Ontology (GO) enrichment analysis of differentially expressed genes was implemented by the GOseq R package, in which gene length bias was corrected. KOBAS software was used to test the statistical enrichment of differential expression genes in KEGG pathways[57, 58]. PPI analysis of differentially expressed genes was based on the STRING database, which predicted Protein-Protein Interactions. The Cufflinks v2.1.1 Reference Annotation Based Transcript (RABT) assembly method was used to construct and identify both known and novel transcripts from TopHat alignment results. Picard-tools v1.96 and samtools v0.1.18 were used to sort, mark duplicated reads and reorder the bam alignment results of each sample. GATK2 (v3.2) software was used to perform SNP calling.

### Expression, purification and functional assay of potential PE-degrading enzymes

To verify the degradation effects of glutathione peroxidase (evm.model.contig_3.359) and laccase (evm.model.contig_8.535) that identified in *Alternaria* sp. FB1, the genes encoding these two proteins were respectively cloned and overexpressed in the *E. coli* cells. First, the intact gene encoding glutathione peroxidase or laccase was amplified from the cDNA template of strain FB1 using the KOD One TM PCR Master Mix (TOYOBO, Japan) with corresponding primers (Supplementary Table S3). The PCR product was purified by using a DNA Gel Extraction Kit (TsingKe, China), and then was cloned in the plasmid pMD19-T simple (TAKARA, Japan). The DNA fragment was digested with *Bam*HI/*Xho*I (Thermo Fisher Scientific, USA), respectively, and ligated into corresponding sites of the expression vector pET28a(+) (Merck, Germany). The recombinant plasmids were transformed into competent cells of *E. coli* BL21(DE3) (TsingKe, China), and transformants were incubated in the Luria-Bertani broth (10 g NaCl, 10 g tryptone and 5 g yeast extract per liter of Milli-Q water) supplemented with 50 μg/mL kanamycin at 37 °C. Protein expression was induced at an OD600 around 0.6 with 0.1 mM isopropyl-1-thio-β-D-galactopyranoside (ITPG), and the cells were cultured for further 20 h at 16 °C. Recombinant proteins were purified with a HisTrapTM HP (GE Healthcare, Sweden) by an AKTA pure system (GE Healthcare, Sweden), and dialyzed against filtered and sterilized seawater for 4 h. The purified proteins were checked by SDS-PAGE, and visualized with Coomassie Bright Blue R250. The degradation effects were detected in a solution containing respective proteins at a final concentration of 0.1 mg/mL with PE films (type ET311350, 0.25mm in thickness) at 30 °C for 48 h. The surface morphology and molecular weight of the PE films were respectively checked by SEM and GPC as described above.

### Data availability

The complete genome sequence of *Alternaria* sp. FB1 has been deposited at GenBank under the accession number PRJNA672824. Raw sequencing reads for transcriptomic analysis have been deposited at NCBI under accession numbers SRR15043810 and SRR15043809. Mass spectrometry analyses of components in 60-day fungal treatment sample at different retention times were shown in Supplementary Figures S6-S12. Mass spectrometry analyses of components in 120-day fungal treatment sample at different retention times were shown in Supplementary Figures S13-S25.

## Acknowledgements

This work is funded by the Major Research Plan of the National Natural Science Foundation (Grant No. 92051107), Key Deployment Projects of Center of Ocean Mega-Science of the Chinese Academy of Sciences (Grant No. COMS2020Q04), National Key R and D Program of China (Grant No. 2018YFC0310800), Strategic Priority Research Program of the Chinese Academy of Sciences (Grant No. XDA22050301), and Taishan Young Scholar Program of Shandong Province (tsqn20161051) for Chaomin Sun.

## Author Contributions

RG and CS conceived and designed the study. RG performed most of experiments. RL helped to purify proteins. RG and CS analyzed the data. RG wrote the manuscript. CS revised the manuscript. All authors read and approved the final manuscript.

