## Supplemental Figures and Tables for "A marine fungus efficiently degrades polyethylene"

### Supplementary Figures

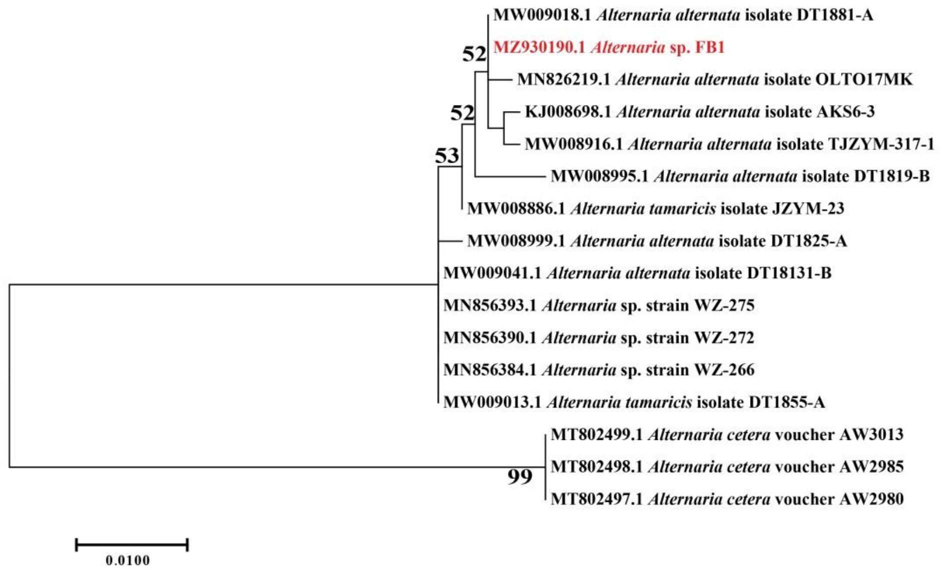


**Figure S1. ITS-based phylogenetic tree of *Alternaria* sp. FB1 with other related fungal strains obtained from the GenBank.** The accession number of each ITS is indicated after the species names. The phylogenetic tree was constructed by the neighbor-joining method and numbers above the branches are bootstrap values based on 1000 replicates. Bar, 0.01 substitutions per nucleotide position.

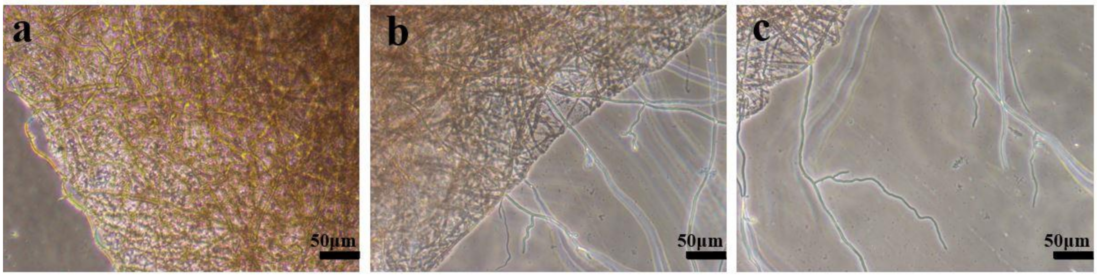


**Figure S2. The light microscopy observation of the mycelium morphology of strain FB1 in the seawater supplemented with or without the PE film for 120 days. a,** The mycelium morphology of strain FB1 incubated in the seawater for 120 days. **b, c,** The mycelium morphology of strain FB1 incubated in the seawater supplemented with the PE film for 120 days.

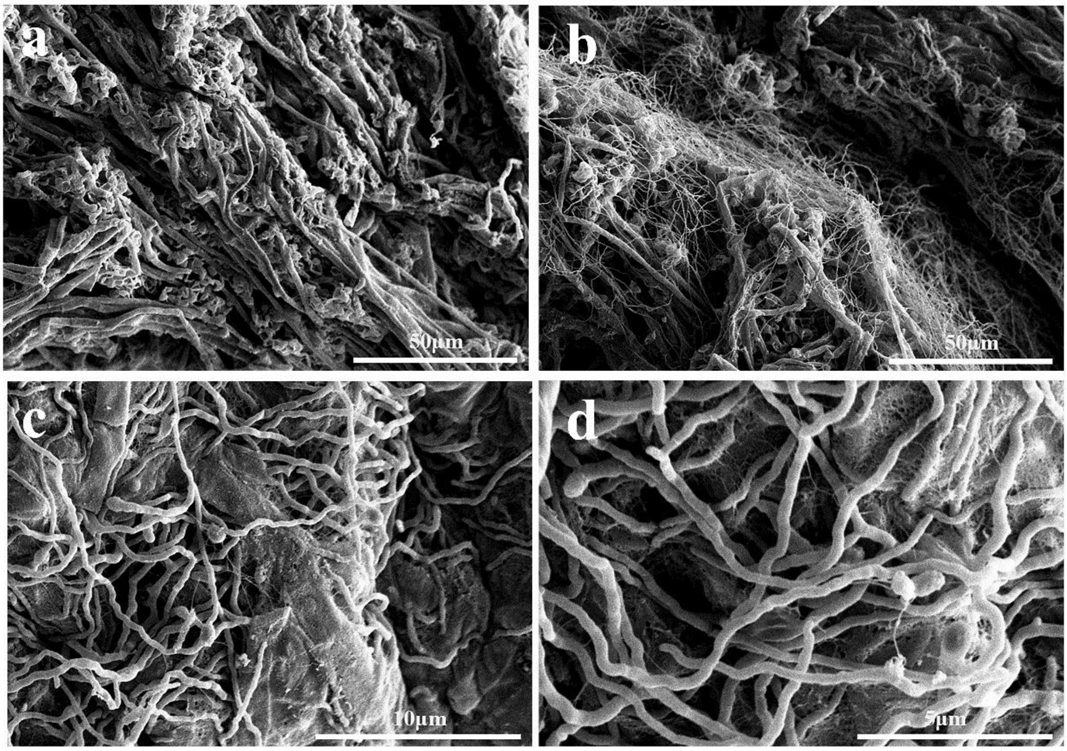


**Figure S3. The SEM observation of the growth of strain FB1 in the seawater supplemented with or without the PE film for 120 days. a**, The SEM observation of the mycelium morphology of strain FB1 in the seawater for 120 days. **b-d**, The SEM observation of mycelium morphology of strain FB1 in the seawater supplemented with the PE film for 120 days.

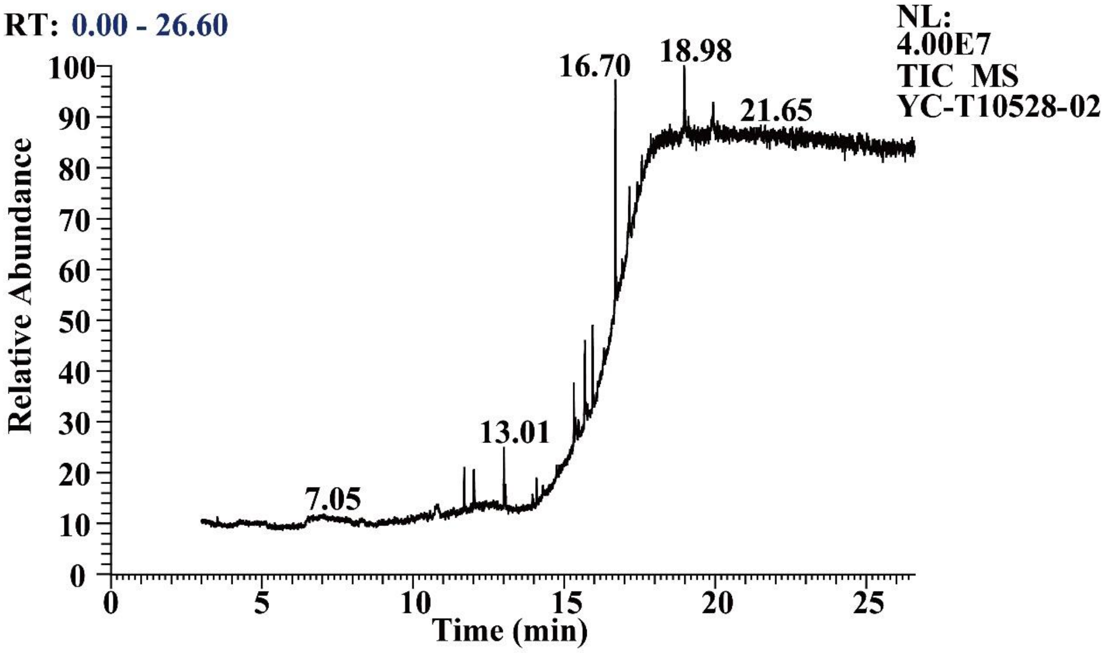


**Figure S4. GC-MS chromatograms of the products extracted from PE biodegradation by *Alternaria* sp. FB1 for 60 days.**

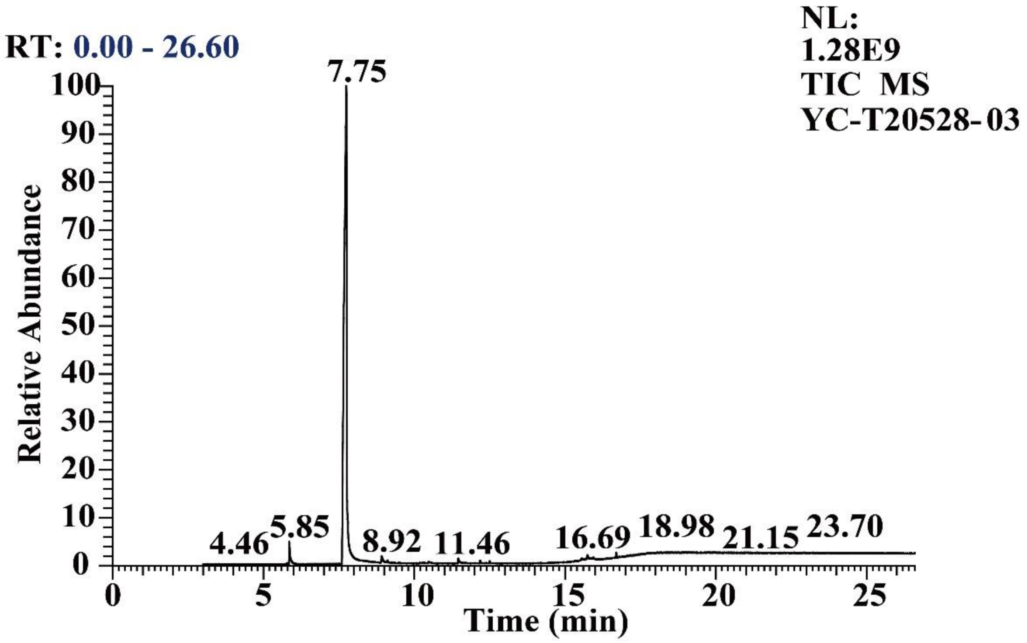


**Figure S5. GC-MS chromatograms of the products extracted from PE biodegradation by *Alternaria* sp. FB1 for 120 days.**

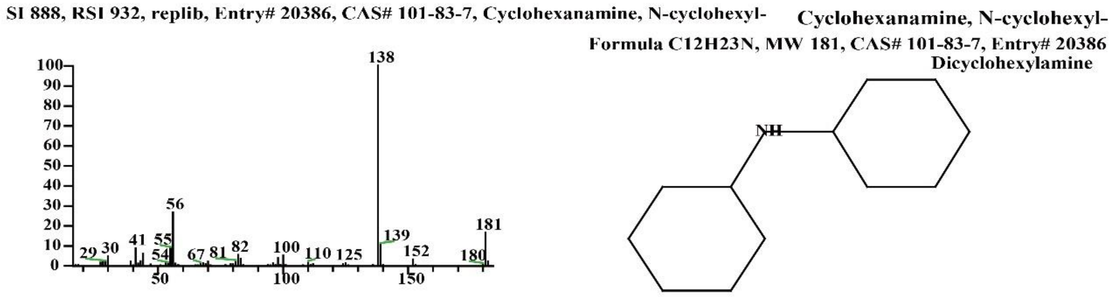


**Figure S6. Mass spectrometry analysis of a component at the retention time of 11.70 min identified as Cyclohexanamine, N-cyclohexyl-.**

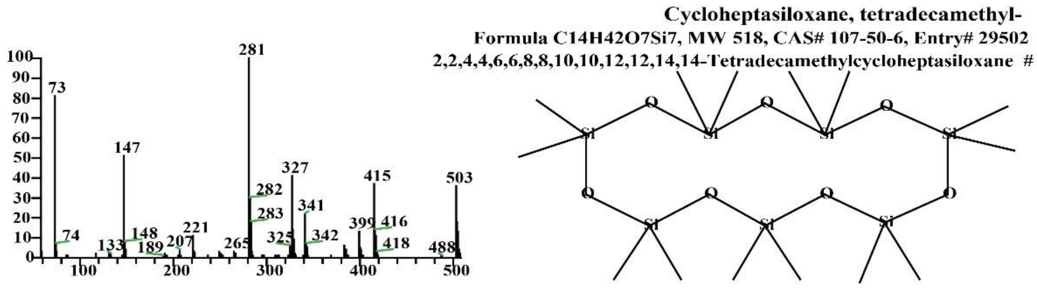


**Figure S7. Mass spectrometry analysis of a component at the retention time of 12.01 min identified as Cycloheptasiloxane, tetradecamethyl-.**

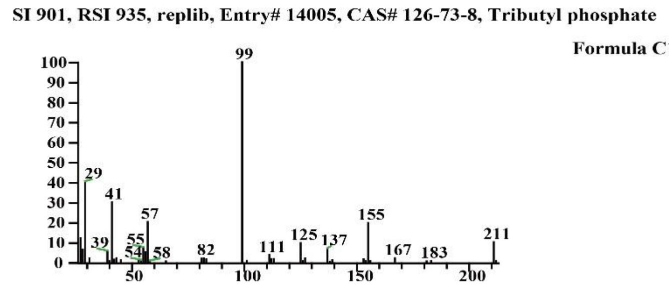


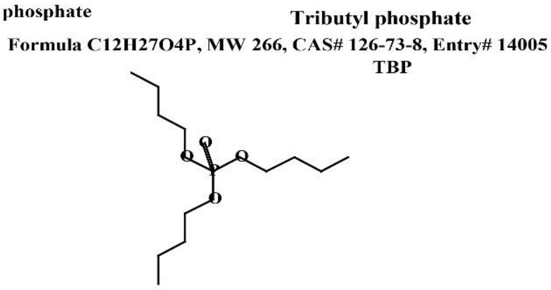


**Figure S8. Mass spectrometry analysis of a component at the retention time of 13.01 min identified as Tributyl phosphate.**

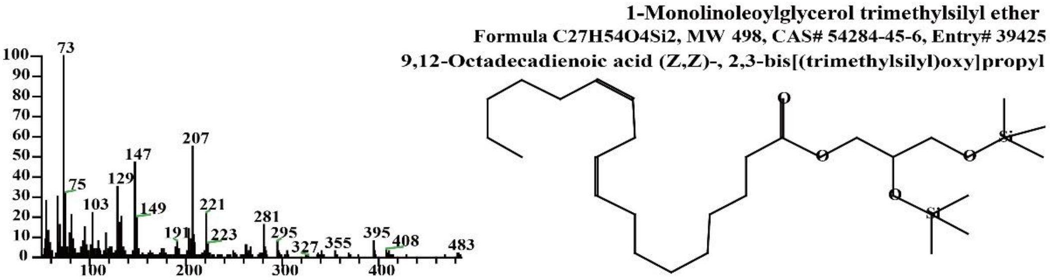


**Figure S9. Mass spectrometry analysis of a component at retention times of 15.32 min, 15.69 min, 15.94 min, 17.58 min identified as 1-Monolinoleoylglycerol trimethylsilyl ether.**

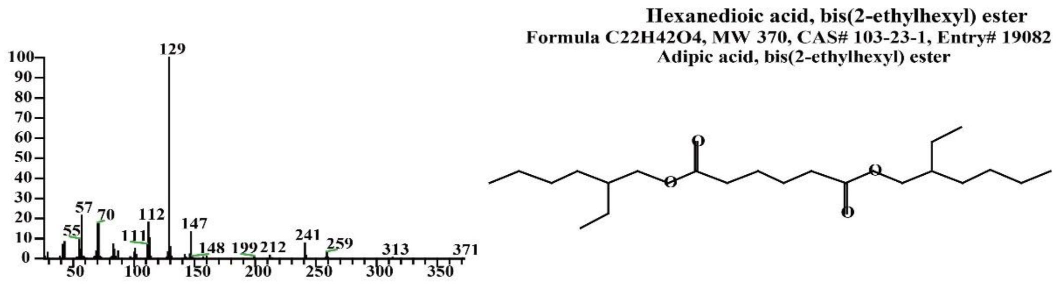


**Figure S10. Mass spectrometry analysis of a component at the retention time of 16.70 min identified as Hexanedioic acid, bis(2-ethylhexyl) ester.**

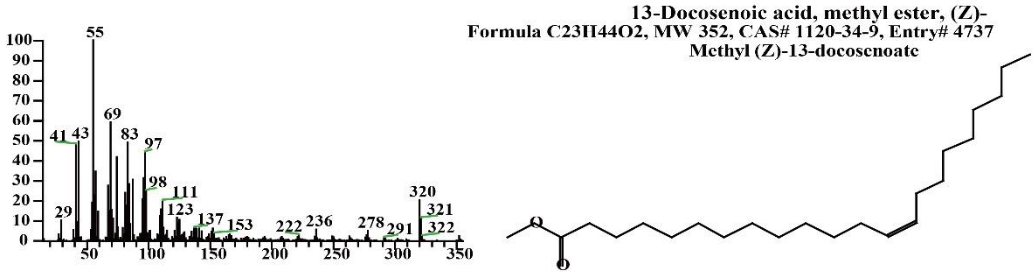


**Figure S11. Mass spectrometry analysis of a component at retention time of 17.16 min identified as13-Docosenoic acid, methyl ester, (Z)-.**

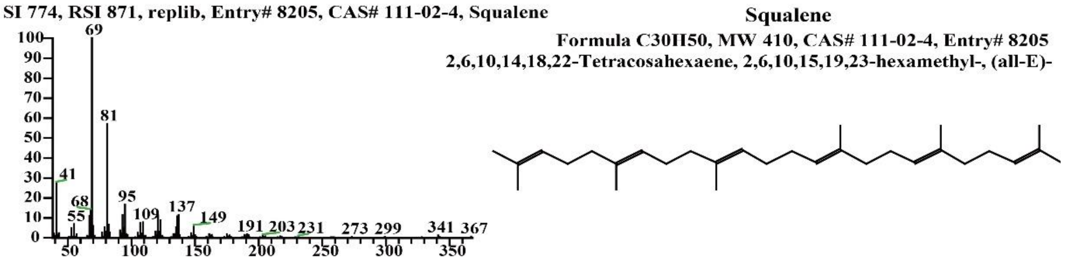


**Figure S12. Mass spectrometry analysis of a component at the retention time of 18.98 min identified as Squalene.**

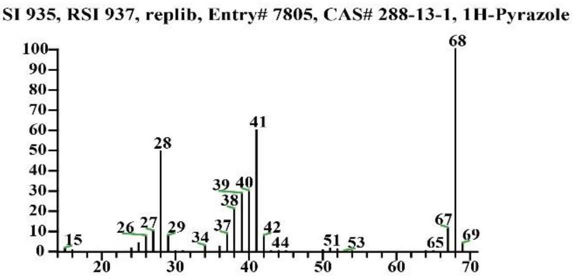


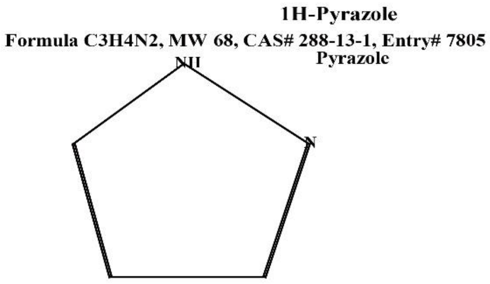


**Figure S13. Mass spectrometry analysis of a component at the retention time of 5.85 min identified as 1H-Pyrazole.**

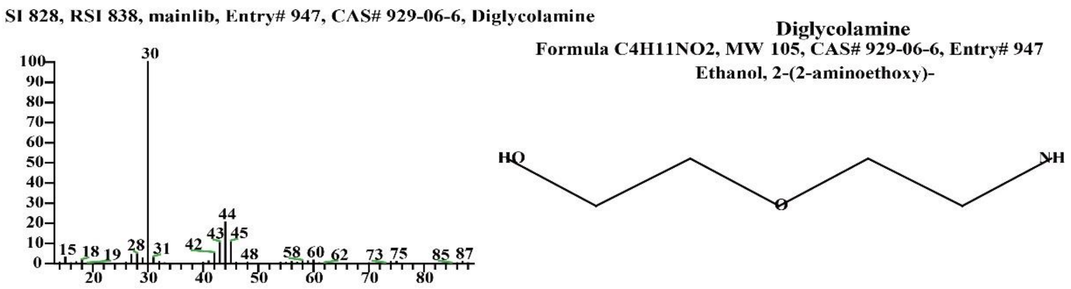


**Figure S14. Mass spectrometry analysis of a component at the retention time of 7.75 min identified as Diglycolamine.**

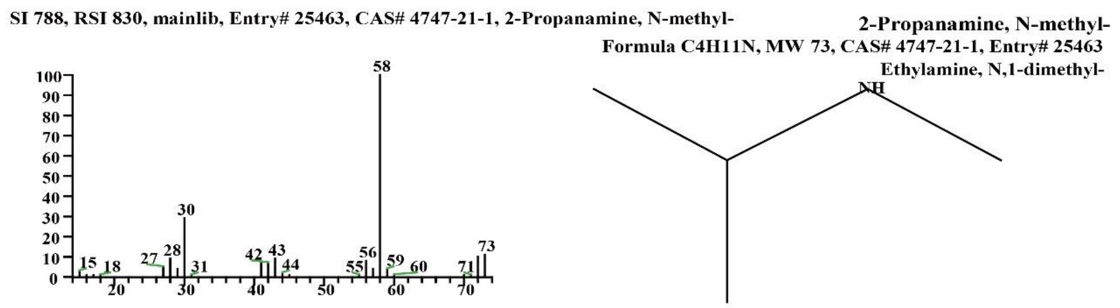


**Figure S15. Mass spectrometry analysis of a component at the retention time of 8.92 min identified as 2-Propanamine, N-methyl-.**

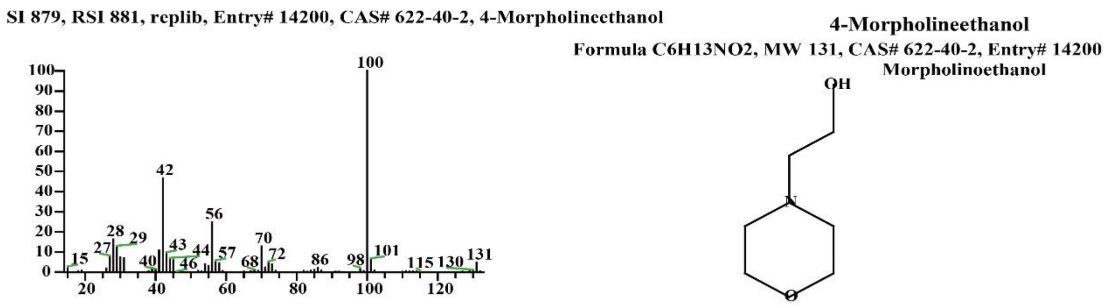


**Figure S16. Mass spectrometry analysis of a component at the retention time of 9.12 min identified as 4-Morpholineethanol.**

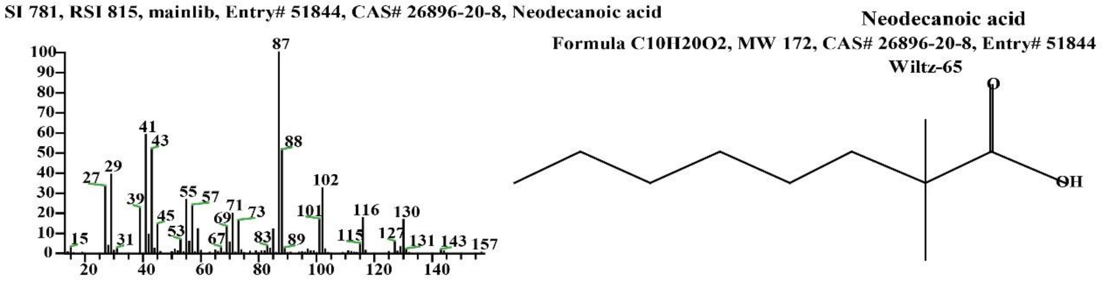


**Figure S17. Mass spectrometry analysis of a component at the retention time of 10.47 min identified as Neodecanoic acid.**

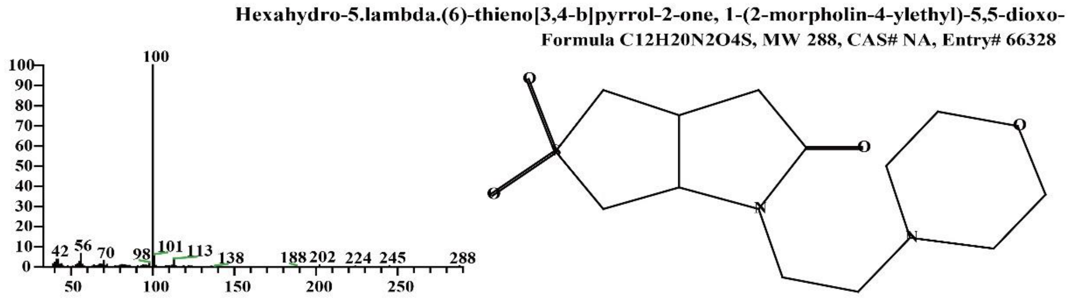


**Figure S18. Mass spectrometry analysis of a component at the retention time of 11.46 min identified as Hexahydro-5.lambda.(6)-thieno[3,4-b]pyrrol-2-one, 1-(2-morpholin-4-ylethyl)-5,5-dioxo-.**

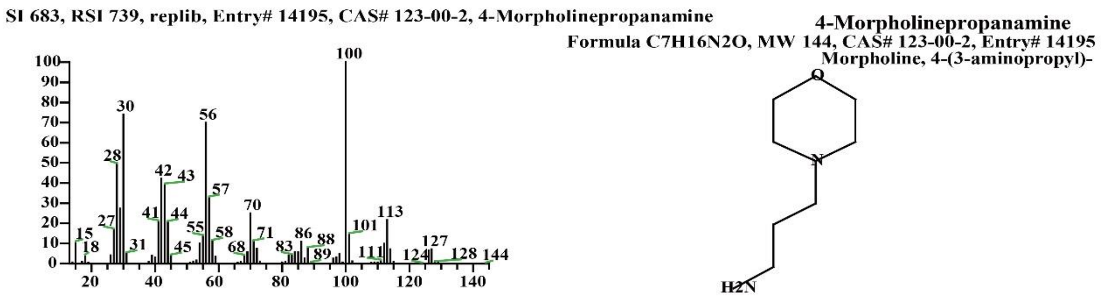


**Figure S19. Mass spectrometry analysis of a component at the retention time of 12.18 min identified as 4-Morpholinepropanamine.**

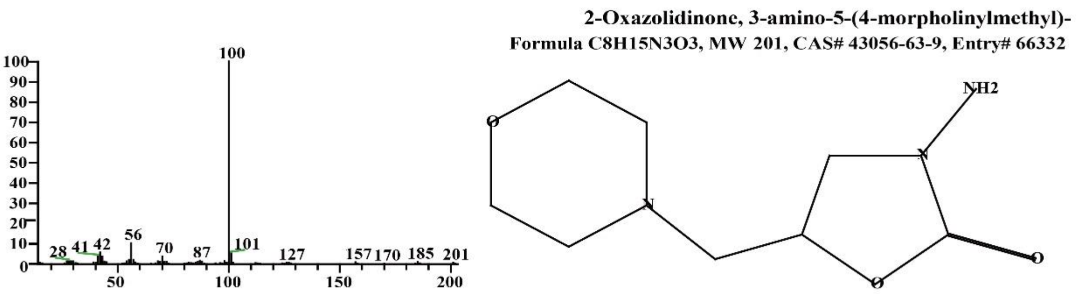


**Figure S20. Mass spectrometry analysis of a component at the retention time of 12.50 min identified as 2-Oxazolidinone, 3-amino-5-(4-morpholinylmethyl)-.**

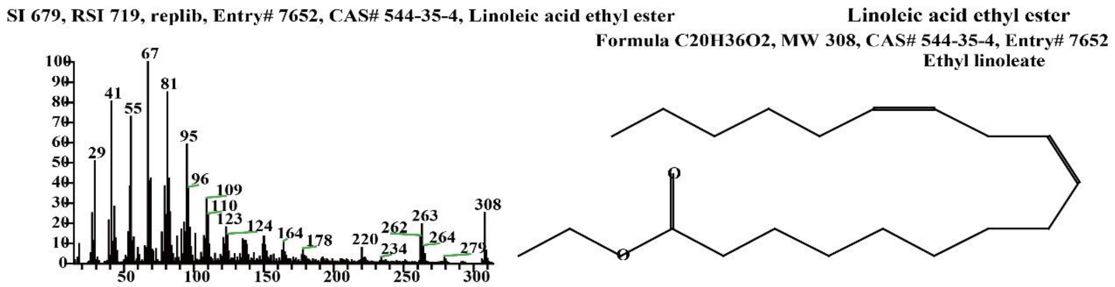


**Figure S21. Mass spectrometry analysis of a component at the retention time of 15.56 min identified as Linoleic acid ethyl ester.**

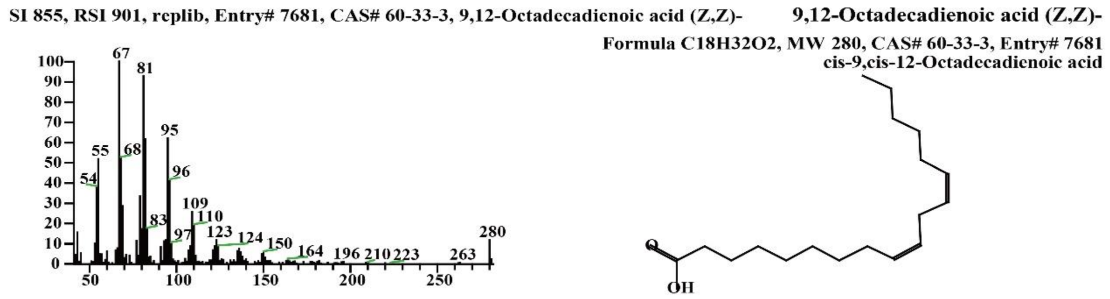


**Figure S22. Mass spectrometry analysis of a component at the retention time of 15.74 min identified as 9,12-Octadecadienoic acid (Z,Z)-.**

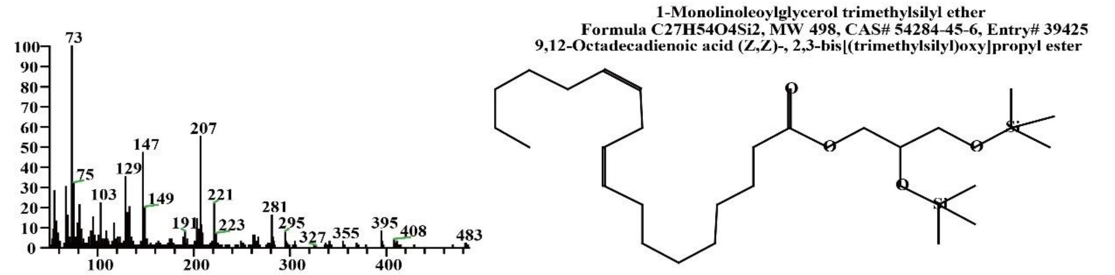


**Figure S23. Mass spectrometry analysis of a component at the retention time of 15.86 min identified as 1-Monolinoleoylglycerol trimethylsilyl ether.**

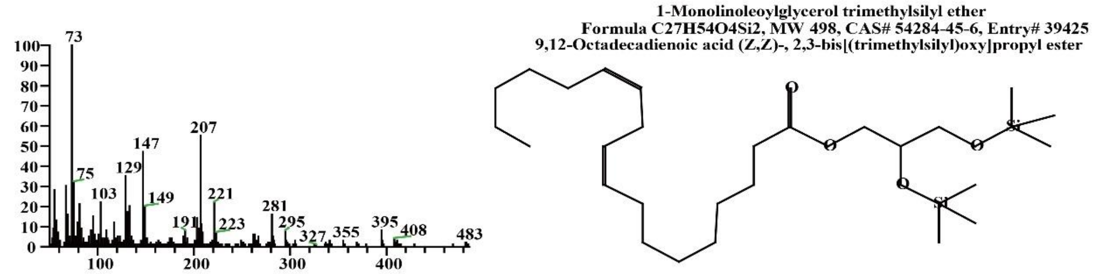


**Figure S24. Mass spectrometry analysis of a component at the retention time of 15.94 min identified as 1,8-Dioxa-5-thiaoctane, 8-(9-borabicyclo[3.3.1]non-9-yl)-3-(9-borabicyclo[3.3.1]non-9-yloxy)-1-phenyl-.**

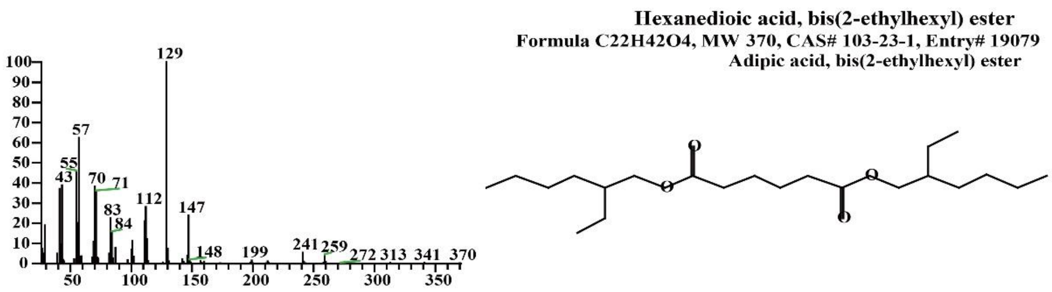


**Figure S25. Mass spectrometry analysis of a component at the retention time of 16.69 min identified as Hexanedioic acid, bis(2-ethylhexyl) ester.**

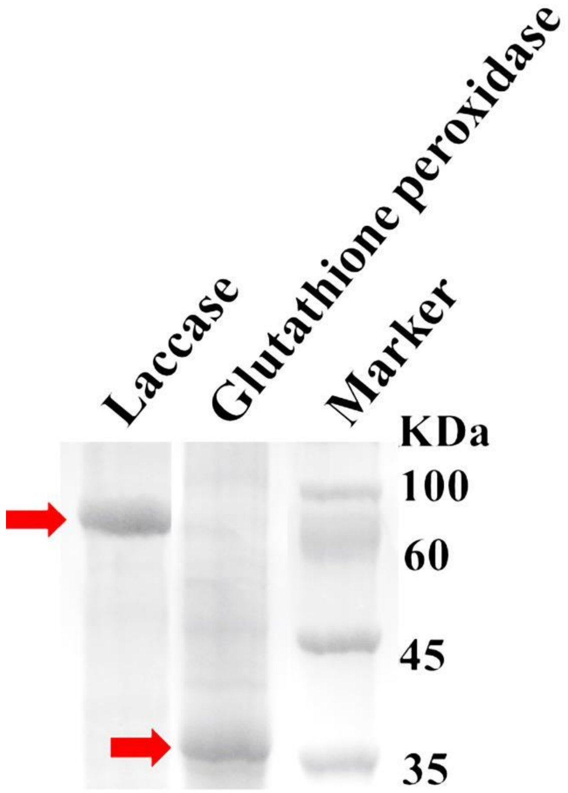


**Figure S26. Coomassie blue-stained SDS gel of purified His-tagged Glutathione peroxidase and Laccase.**

**Supplementary Table S1. Mass spectrometric identification of compositions released from the PE film treated by *Alternaria* sp. FB1 for 60 days.**

| Retention time | Peak Area | Area % | chemical formula | Compound Name | Cas # |
|---|---|---|---|---|---|
| 11.70 | 3803383.72 | 2.33 | $C_{12}H_{23}N$ | Cyclohexanamine, N-cyclohexyl- | 101-83-7 |
| 12.01 | 5640482.26 | 3.45 | $C_{14}H_{42}O_7Si_7$ | Cycloheptasiloxane, tetradecamethyl- | 107-50-6 |
| 13.01 | 7789352.35 | 4.77 | $C_{12}H_{27}$ | Tributyl phosphate | 126-73-8 |
| 15.32 | 8879994.65 | 5.44 | $C_{27}H_{54}O_4Si_2$ | 1-Monolinoleoylglycerol trimethylsilyl ether | 54284-45-6 |
| 15.69 | 14727808.96 | 9.02 | $C_{27}H_{54}O_4Si_2$ | 1-Monolinoleoylglycerol trimethylsilyl ether | 54284-45-6 |
| 15.94 | 7910509.73 | 4.84 | $C_{27}H_{54}O_4Si_2$ | 1-Monolinoleoylglycerol trimethylsilyl ether | 54284-45-6 |
| 16.70 | 26809165.70 | 16.42 | $C_{22}H_{42}O_4$ | Hexanedioic acid, bis(2-ethylhexyl) ester | 103-23-1 |
| 17.16 | 12894410.05 | 7.90 | $C_{23}H_{44}O_2$ | 13-Docosenoic acid, methyl ester, (Z)- | 1120-34-9 |
| 17.58 | 52152219.34 | 31.94 | $C_{27}H_{54}O_4Si_2$ | 1-Monolinoleoylglycerol trimethylsilyl ether | 54284-45-6 |
| 18.98 | 22675990.39 | 13.89 | $C_{30}H_{50}$ | Squalene | 111-02-4 |

**Supplementary Table S2. Mass spectrometric identification of compositions released from the PE film treated by *Alternaria* sp. FB1 for 120 days.**

| Retention time | Peak Area | Area % | chemical formula | Compound Name | Cas # |
|---|---|---|---|---|---|
| 5.85 | 160827301.91 | 1.98 | $C_3H_4N_2$ | 1H-Pyrazole | 288-13-1 |
| 7.75 | 7586192268.64 | 93.28 | $C_4H_{11}NO_2$ | Diglycolamine | 929-06-6 |
| 8.92 | 53491610.03 | 0.66 | $C_4H_{11}N$ | 2-Propanamine, N-methyl- | 4747-21-1 |
| 9.12 | 8303541.90 | 0.10 | $C_6H_{13}NO_2$ | 4-Morpholineethanol | 622-40-2 |
| 10.47 | 17177624.61 | 0.21 | $C_{10}H_{20}O_2$ | Neodecanoic acid | 26896-20-8 |
| 11.46 | 40745877.94 | 0.50 | $C_{12}H_{20}N_2O_4S$ | Hexahydro-5.lambda.(6)-thieno[3,4-b]pyrrol-2-one, 1-(2-morpholin-4-ylethyl)-5,5-dioxo- | NA |
| 12.18 | 10690688.81 | 0.13 | $C_7H_{16}N_2O$ | 4-Morpholinepropanamine | 123-00-2 |
| 12.50 | 6114688.36 | 0.08 | $C_8H_{15}N_3O_3$ | 2-Oxazolidinone, 3-amino-5-(4-morpholinylmethyl)- | 43056-63-9 |
| 15.56 | 23251808.47 | 0.29 | $C_{20}H_{36}O_2$ | Linoleic acid ethyl ester | 544-35-4 |
| 15.74 | 54484769.63 | 0.67 | $C_{18}H_{32}O_2$ | 9,12-Octadecadienoic acid (Z,Z)- | 60-33-3 |
| 15.86 | 9971750.86 | 0.12 | $C_{27}H_{54}O_4Si_2$ | 1-Monolinoleoylglycerol trimethylsilyl ether | 54284-45-6 |
| 15.94 | 16603906.65 | 0.20 | $C_{27}H_{54}O_4Si_2$ | 1-Monolinoleoylglycerol trimethylsilyl ether | 54284-45-6 |
| 16.69 | 15787407.31 | 0.19 | $C_{22}H_{42}O_4$ | Hexanedioic acid, bis(2-ethylhexyl) ester | 103-23-1 |

**Supplementary Table S3. Primers used for construction of vectors for expression of potential PE-degrading enzymes.**

| Primer | Sequence |
|---|---|
| Glutathione peroxidase F | GGATCCATGATTCGATTGGGAGGGG |
| Glutathione peroxidase R | CTCGAGTTTGCCGCCCAGCTCCTTC |
| Laccase F | GGATCCATGAGCGAGCACACTGATTCGC |
| Laccase R | CTCGAGGATTGTGTCCGTGTCTAGAATTCTT |
